# A better connected brain relates to lower dogmatism in young and older adults

**DOI:** 10.64898/2026.06.29.735242

**Authors:** Geraldine Rodríguez-Nieto, Stephan P. Swinnen

**Affiliations:** Movement Control and Neuroplasticity Research Group, Biomedical Sciences, KU Leuven, 3001 Leuven, Belgium; KU Leuven Brain Institute, KU Leuven, Belgium; REVAL – Rehabilitation Research Center, Faculty of Rehabilitation Sciences, University of Hasselt, Diepenbeek, Belgium

**Author notes:** **Correspondence: Geraldine Rodriguez Nieto**.

**Keywords:** cognitive flexibility, dogmatism, aging, structural connectivity, DWI, network

## Abstract

BACKGROUND: Cognitive flexibility represents a crucial function in adapting to new environments. In this study we examined the ecological validity of a cognitive flexibility task by studying its relationship with individual traits (dogmatism, dependence on routines and perspective taking). Second, we investigated whether global and local structural brain connectivity properties were related to cognitive flexibility as well as associated traits and their possible age-related differences. METHOD: Thirty-eight young (18-35 years) and thirty-seven older (60-85 years) healthy participants took part in an MRI protocol including a Diffusion Weighted Imaging (DWI) sequence. Participants also performed a Rule-Switching task to measure cognitive flexibility performance and filled in questionnaires assessing dogmatism, dependence on routines and perspective taking. RESULTS: A higher cognitive flexibility was related to lower dogmatism and lower dependence on routines only in young adults. In relation to structural connectivity, we found that: a) global and local connectivity properties negatively predicted dogmatism levels in the full sample, b) local connectivity properties of the inferior frontal gyrus (IFG) positively predicted performance in cognitive flexibility performance in the full sample and in older adults, and c) connectivity between left inferior parietal lobule (IPL) and left putamen negatively predicted dogmatism in older adults. DISCUSSION: A deeper understanding of the shaping of structural networks supports a better understanding of cognitive flexibility and dogmatism in a highly dynamic world.

## 1. INTRODUCTION

Cognitive flexibility is an executive function that refers to the capacity to switch among different mental schemes and, therefore, provides an evolutionary advantage in managing adaptability to new environments. In a constantly changing world faced with new technologies, and altered socio-political environments and climate, cognitive flexibility may support adaptation to new challenges.

Typically, cognitive flexibility has been investigated through neuropsychological tasks in experimental settings and clinical practice. Nonetheless, the ecological validity of these tasks is often neglected. Accordingly, to what extent such (computer-based) tasks relate to daily life remains elusive. Additionally, the impact of cognitive flexibility in daily life may be conceived in different ways. For instance, lack of cognitive flexibility may be reflected in rigidity affecting an individual’s way of thinking (dogmatism), being unable to consider other perspectives, or showing rigidity in daily habits. Evidence from studies in autistic individuals suggest that cognitive flexibility is a heterogeneous construct (Geurts et al. 2009); however evidence from neurotypical individuals is largely missing.

Whereas older adults frequently show a poorer performance on cognitive flexibility tasks (Rodriguez-Nieto et al. 2024; Seer et al., 2022), less is known about differences in cognitive flexibility as assessed through other methods, such as self-reports. Likewise, as executive functions (inhibition, switching and updating) relate to each other in different ways across the lifespan, possibly due to brain structural changes (Diamond, 2013; Lee et al., 2013; Karr et al., 2018; Glisky et al. 2021) it is also possible that the unity-segmentation of processes related to cognitive flexibility vary with aging.

The de-differentiation of cognitive processes with aging may be related to changes in brain structure. As cognitive flexibility refers to the ability to switch among different mental schemes, and, to some extent implies the ability to recruit different cognitive processes (ie. during multitasking or in applying a new approach to an old problem), it may be expected that the features of brain structural connectivity are highly relevant for this domain.

In older as compared to young adults important differences in structural connectivity emerge, such as decreased connectivity strength and efficiency (Coelho et al., 2020; Damoiseaux et al., 2017; Zhao et al., 2015). Further studies have shown that changes in efficiency play a mediatory role in the age-related decline of executive functions (Li et al., 2020; Rodriguez-Nieto et al. 2025). However, no associations were observed between efficiency measures and a Switching (cognitive flexibility) factor (Rodriguez-Nieto et al., 2025). Moreover, to our knowledge, the relationship between structural connectivity and traits associated with cognitive flexibility has not been studied.

The left inferior frontal gyrus (IFG) and the left inferior parietal lobe (IPL) have shown to be commonly active in different types of cognitive flexibility paradigms and are considered to be flexible hubs in the brain (Yeo et al., 2015; Camilleri et al., 2018). In addition, the IFG has consistently shown differences related to executive functioning as an effect of aging and its hypometabolism relates to executive deficits in different neurodegenerative disorders (Schroeter et al., 2012; Heckner et al., 2021). For this reason, we hypothesize that besides the global structural connectivity properties of the brain, the structural connectivity of these specific regions is relevant in cognitive flexibility and associated traits.

Finally, different studies have highlighted the role of the basal ganglia in cognitive flexibility (van Schouwenburg; Uddin, 2021). In our previous functional connectivity study, we observed that a stronger connectivity between the left IPL and the bilateral putamen during the performance of a cognitive flexibility task positively predicted the quality of performance in young individuals. However, in older adults, a stronger left IPL-putamen and left IPL-precuneus connectivity related to a poorer performance (Rodriguez Nieto et al., 2026).

In this study we aimed to investigate: a) the ecological validity of a cognitive flexibility task by determining its relation to associated traits in young and older adults, and b) age differences in traits presumably related to cognitive flexibility (self-reports), hypothesizing that older adults would report lower cognitive flexibility in their daily lives. Given the relationship between structural connectivity efficiency and executive functions -but not with Switching-(Heckner et al. 2021; Rodriguez-Nieto et al. 2025) in older adults, we aimed to: c) test the relationship between structural network connectivity properties and cognitive flexibility, measured with a task not used in our previous study and through self-reports (cognitive flexibility associated traits) in young and older adults. In addition, given the crucial role of the left inferior frontal cortex (IFC) and the left inferior parietal lobe (IPL) in cognitive flexibility (Rodriguez-Nieto et al., 2022; Worringer et al. 2019) and of the role of the functional connectivity of the precuneus and putamen with the left IPL in cognitive flexibility performance (Rodriguez-Nieto et al., 2026), we tested whether cognitive flexibility performance and related traits are associated to the structural connectivity properties of the IFC and IPL regions (d), and to the structural connectivity between the IPL and precuneus and putamen (e).

## 2. METHOD

### 2.1 Participants

Thirty-eight young (YA: mean age=23.38 SD=3.81, range=18-35 years old, 25 female) and thirty-seven older (OA: mean ± SD age=69.52, SD=5.42, range=60-85 years old, 20 female) healthy participants were recruited for this study. All participants reported absence of neurological and psychiatric disorders. The Montreal Cognitive Assessment (MoCA) was used to exclude participants with mild cognitive impairment (YA: mean score=27.4, range=24-30; OA: mean score=27.1, range=22-30). One older participant scored below the cutoff point (score=22; cutoff=24: Islam et al., 2023) and was excluded for that reason. The handedness of participants was assessed with the Edinburgh Handedness questionnaire. The handedness proportion of participants was representative of that reported for the general population (Willems et al., 2014): 37 young adults and 37 older adults were right-handed; three younger adults and two older adults were left-handed, and one older adult was ambidextrous. The experiment was approved by the local ethical committee of KU Leuven (Protocol ID: s61577) and all participants gave written informed consent.

### 2.2 Procedure

Participants took part in two sessions. In the first session, participants filled in self-reports (including questionnaires to assess cognitive flexibility related traits) and performed computer tasks. During the second session, participants took part in an MRI protocol.

Prior to entering the MRI room, participants performed 16 practice trials of the Rule Switching task used to assess cognitive flexibility. During the MRI session, they performed the task during fMRI data collection. The behavioral data of this scan was used for the present study. Magnetic Resonance Spectroscopy (MRS), Diffusion Weighted Imaging (DWI) and resting state Magnetic Resonance Imaging (rsMRI) were also collected. For this study, we only investigated the output from the DWI data.

### 2.3 Behavioral task

In the Rule-switching flexibility task (adapted from Sekutowicz et al., 2016) participants utilized a button box to respond to stimuli displayed on the screen. They indicated whether a displayed number was lower or higher than five (left or right button press) when the number was preceded by a cue and simultaneously enclosed by a circle. Additionally, they determined whether the number was odd or even (left or right button press) when preceded by a cue and enclosed by a square (see Figure 1). The task included two types of trials: Repeat trials, where the rule remained the same as the previous trial, and Switch trials, where the rule changed. The sequence of trials was pseudo-randomized. The number of correct answers in the Repeat trials was subtracted from the number of correct answers in the Switch trials, which constituted the cognitive flexibility index (a higher index indicated higher cognitive flexibility).

**Figure 1.**
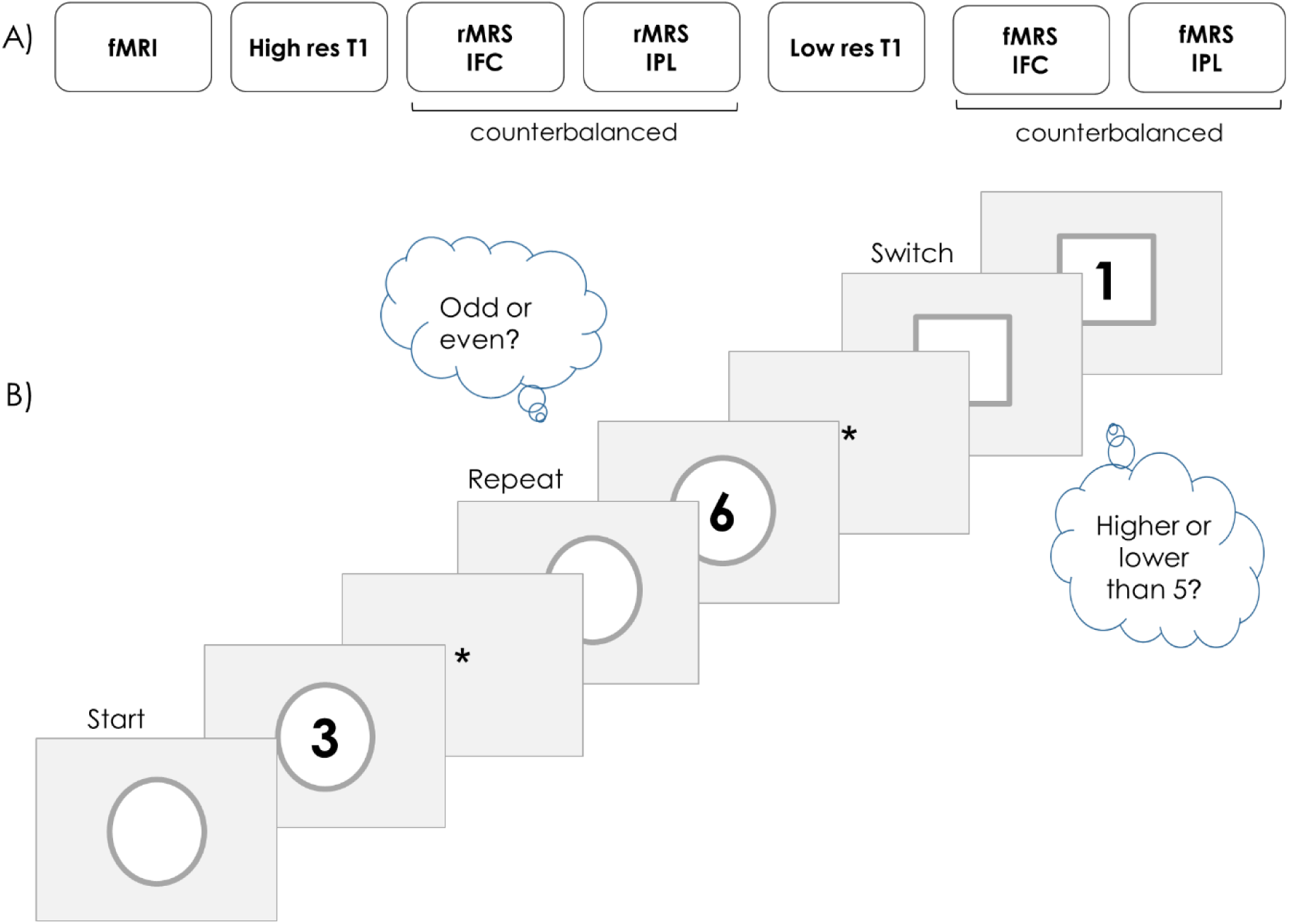
A) MRI session protocol. rMRS: resting state MRS; fMRS: MRS during cognitive flexibility performance; B) Cognitive flexibility task design. Individuals followed one of two rules (decide if the number was odd or even, or decide whether the number was higher or lower than 5) according to the shape that surrounded the number. If the rule was the same as in the previous trial, this was a Repeat trial, whereas if the rule was different from the previous trial, this was a Switch trial. The performance on Switch and Repeat trials was compared to generate a cognitive flexibility index.

During the fMRI session, participants completed 94 trials, with a cue duration of 300ms, cue+stimulus duration of 1700ms, and inter-trial interval (ITI) ranging from 3 to 4 seconds. The task was implemented using PsychoPy (Peirce, 2007). Five datapoints of young participants for this task were excluded due to technical problems that led to recording errors.

### 2.4 Cognitive flexibility associated traits

#### Dogmatism scale (Shearmann and Levine, 2006)

This type of dogmatism is described as a cognitive style characterized by close-mindedness to different opinions or perspectives. The updated dogmatism scale is correlated positively with social dominance (need of control over others) and negatively with perspective-taking and empathetic concern (Shearman and Levine, 2006). The output of this self-report also holds a positive correlation with anxiety scores extracted from the Beck Depression Inventory (Rodriguez-Nieto et al., 2025). The scale is constituted by eleven items in a five-point Likert scale.

#### DOG scale (Crowson et al., 2015)

This type of dogmatism is defined as an unchangeable and unjustified certainty in one’s beliefs (eg. being absolutely sure about the ‘big truths’ in life). This scale is correlated with authoritarianism and zealotry (Altemeyer 2002). The scale is constituted by twenty-two items in a 5-point Likert scale.

#### Dependence on Routines (Zmigrod et al., 2018)

This scale evaluates dependence on daily routines and uneasiness or intolerance when these routines are disrupted. The questionnaire comprises eight items evaluated on a seven-point Likert scale ranging from 1 to 5 (i.e., Not at all characteristic of me - Entirely characteristic of me).

#### Perspective taking (subscale from the Interpersonal Reactivity Index; Davis, 1983)

This subscale measures an individual’s self-reported propensity to spontaneously engage in mental-state inference (adopt the standpoint of other). It is comprised of seven items in a five-point Likert scale (i.e., Does not describe me well – Describes me very well).

### 2.5 Imaging data acquisition

MRI data was acquired with a Philips Achieva 3T scanner equipped with a 32-channel head coil. A high-resolution three-dimensional T1-weighted image was obtained using the MPRAGE sequence (time repetition/time echo = 9.6 ms/4.6 ms; voxel size = 0.98 mm × 0.98 mm × 1.2 mm; field of view = 192 × 250 × 250; 160 coronal slices; scan time approximately 7 minutes).

Diffusion MRI data were acquired using a single-shot echo planar imaging sequence with the following parameters: dMRI volumes with b-values = 700 s/mm2 (16 gradient directions), 1,200 s/mm2 (30 gradient directions), and 2,800 s/mm2 (50 gradient directions); 6 interspersed volumes without diffusion weighting (b=0 s/mm2); flip angle=90◦; phase encoding direction = posterior to anterior (PA); field of view = 240 × 240 × 140 mm3; voxel size = 2.5 × 2.5 × 2.5 mm3, TE/TR =74/5,000 ms; multiband factor = 2; SENSE = 2; matrix size = 96 × 96; 56 transverse slices; total scan time=∼9 min. We also acquired five b = 0 s/mm2 images with reversed phase encoding (AP) for the purpose of susceptibility-induced distortion correction.

### 2.6 MRI processing

The MRtrix3 (Tournier et al., 2019) standard structural connectome construction pipeline (Smith and Connelly, 2019; available at https://github.com/BIDS-Apps/MRtrix3_connectome) was applied to dMRI and T1W data. Where necessary, this pipeline also incorporates commands from FSL (Jenkinson et al., 2012) and Freesurfer (Fischl, 2012) software packages. Brain parcellation was performed according to the Desikan atlas (Desikan et al., 2006). In brief, dMRI data were denoised (Veraart et al., 2016), Gibbs unringed (Kellner et al., 2016), and corrected for eddy current distortions, motion, and susceptibility induced distortions (Andersson et al., 2003, 2016, 2017; Andersson and Sotiropoulos, 2016). Three-tissue response functions representing single-fiber white matter, gray matter and cerebrospinal fluid were obtained from the corrected dMRI data using an unsupervised approach (Dhollander et al., 2016). Three-tissue constrained spherical deconvolution (CSD) was performed for each participant, using the averaged (across all participants) response functions for each tissue type with the multi-shell multi-tissue CSD algorithm (Jeurissen et al., 2014), resulting in the white matter fiber orientation distribution (FOD) for each voxel. Joint bias field correction and global intensity normalization of the 3-tissue parameters was performed in the log-domain (Dhollander et al., 2021). Subject’s T1W image was also registered to the mean b = 0 s/mm2 (corrected) image via rigid-body transformation (Bhushan et al., 2015).

Following the initial processing, tractograms were generated. Thus, for each participant, the 2nd-order integration over FODs algorithm (iFOD2; Tournier et al., 2010) and the anatomically constrained tractography (ACT; Smith et al., 2012) with dynamic seeding (Smith et al., 2015a), FOD amplitude threshold 0.06, step size of 1.25 mm, length of 5–250 mm, and backtracking (Smith et al., 2012) were used to generate 10 million probabilistic streamlines. Furthermore, each streamline was assigned a weight, computed using the spherical-deconvolution informed filtering of tractograms (SIFT2; see Smith et al., 2015a). Based on each participant’s tractogram, an individual connectome was computed using 84 regions-of-interest parcellated in native space [cortex and cerebellum: Dale et al. (1999); Desikan et al. (2006); subcortical regions: Patenaude et al. (2011); see Smith et al. (2015a)], with connection strengths calculated by summing the weights of the relevant streamlines scaled by the proportionality coefficient (Smith et al., 2015a). These 84 nodes were used for further analyses. The DWI datapoints from one young adult and two older adults were excluded due to technical problems leading to non-usable data.

### 2.7 Efficiency, Clustering and Connection Strength

Figure 2 summarizes the procedure for processing the structural connectivity data. In sum, after the pre-processing of the DWI output and the generation of the connectome matrices, graph theory network analyses were conducted to extract global and local connectivity properties. The Brain Connectivity Toolbox (Rubinov and Sporns, 2010), implemented in MATLAB (TheMathWorks Inc., Natick, MA), was used to compute weighted, undirected network metrics, in particular: global efficiency, local efficiency, clustering and strength. Global efficiency is defined as the average inverse shortest path length (Latora and Marchiori, 2001), indicating the integration of the entire network (the ability to combine specialized information from distributed regions; Rubinov and Sporns, 2010). The local efficiency of a node in the graph is the average global efficiency of the subgraph induced by the neighbors of the node (Latora and Marchiori, 2001). The average local efficiency is the average of the local efficiencies of each node in the network and reflects local network integration. Clustering is a measure of segregation (the ability of specialized processing within densely interconnected modules -or clusters-; Rubinov and Sporns, 2010) and it is defined as the proportion of connected edges in the local neighboring sub-graph for each node. Finally, strength refers to the weight of the edge connecting two nodes (in tractography, it reflects the number of reconstructed streamlines linking pairs of brain regions; Fornito et al., 2016); the strength of the entire network is the average of the strength of the edges in the network.

**Figure 2.**
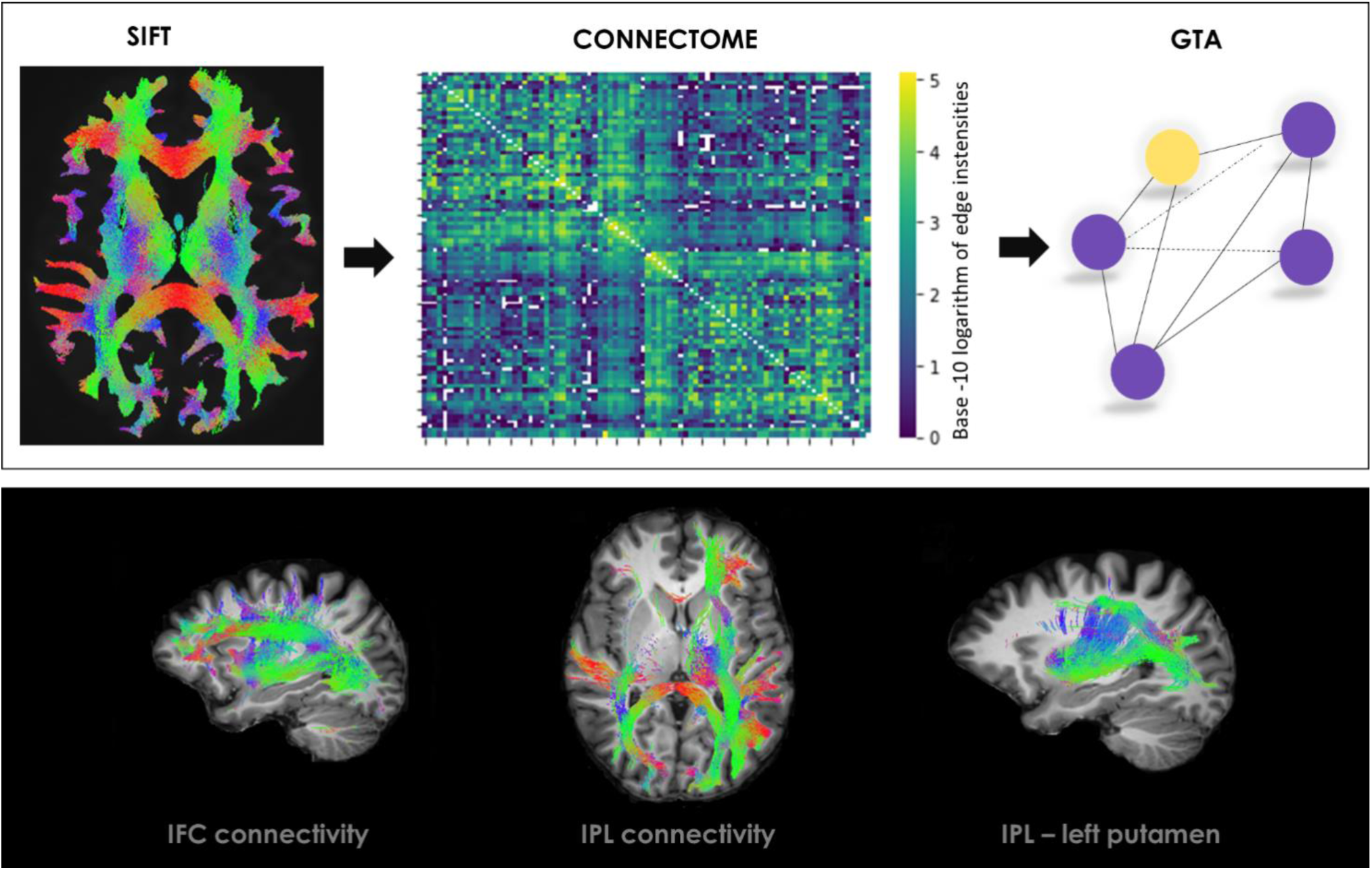
Top panel: Left: Illustrative tractogram reconstruction (SIFT2) obtained after preprocessing; Middle: In combination with the parcellation information a connectivity matrix with 84 nodes (brain regions) is created; Right: Graph Theory Analysis (GTA) is applied to the connectivity matrix to extract network features. Bottom panel: Connectivity of the left inferior frontal cortex, the inferior parietal lobe and inferior parietal lobe-putamen from one participant is shown for illustrative purposes.

The efficiency, clustering and strength indices of the inferior frontal gyrus pars triangularis (IFJpt) and the IPL were also extracted, as these regions are crucial nodes in the processing of cognitive flexibility (Rodriguez Nieto et al., 2025). Finally, the connectivity strength indices of the left IPL with the left putamen, the right putamen and the precuneus were also extracted, as we previously reported that the functional connectivity between these regions is predictive of cognitive flexibility performance (Rodríguez Nieto et al., 2026).

### 2.8 Statistical Analyses

The data was inspected to check for normality and outliers. One datapoint from one young participant was excluded for being above 3.5 standard deviations from the mean in the DOG scale.

T-tests were conducted to examine age differences in cognitive flexibility, associated traits and structural connectivity properties. When the data did not show a normal distribution (Dependence on Routines and Perspective taking), U-Mann Whitney tests were conducted. Multiple comparisons correction (with False Discovery Rate, Benjamini-Hochberg) was applied.

Correlations were used to investigate the relationship between cognitive flexibility (Rule Switching task) and associated traits (dogmatism, dependence on routines and perspective taking). These analyses were performed in young and old adults separately in view of the previously reported differences in the cognitive flexibility task (Rodriguez-Nieto et al., 2024). When the data did not show a normal distribution, Spearman correlations were applied. Multiple comparisons correction with the Benjamini-Hochberg method was applied.

To investigate whether global brain connectivity properties (global efficiency, strength and clustering) and their interaction with age (in years) could predict cognitive flexibility and related traits (dogmatism, dependence on routines and perspective taking), linear regression models were conducted (one model per variable: cognitive flexibility task, dogmatism -one per each self-report-, dependence on routines and perspective taking). Local efficiency was excluded from these models given its high correlation with clustering (YA: r=.98 <.001 and OA: r=.99 <.001).

Further regression analyses were conducted to investigate whether clustering and connectivity strength of IFG and IPL and their interaction with age (in years) could predict cognitive flexibility and associated traits (one model per variable). Efficiency from these regions was not included given its high correlation with clustering (r=.99 <.001 for all cases).

Finally, regression models were used to examine whether the strength of the connectivity between the left IPL and the left putamen, right putamen, left precuneus and right precuneus, and their interaction with age (in years) predicted cognitive flexibility and associated traits. This selection was based on our previous findings showing a relationship between functional connectivity in these pairs of nodes and the execution of cognitive flexibility performance (Rodríguez-Nieto et al., 2026).

## 3. RESULTS

### 3.1 Age differences in cognitive flexibility, associated traits and structural connectivity properties

The descriptive statistics and independent sample tests according to age are displayed in Table 1. In sum, young individuals showed a better performance in the Rule Switching task but no age differences were observed in either of the cognitive flexibility associated traits (dogmatism, dependence on routines and perspective taking). Young individuals showed higher levels of global and local efficiency, clustering and strength than older adults. Likewise, young adults showed higher levels of efficiency, clustering and strength in the IPL and in the IFGpt and in the connectivity strength of the left IPL with the left putamen and with the right precuneus. Interestingly, older adults showed a stronger connectivity between of left IPL with the right putamen and with the left precuneus, but this difference was not significant.

**Table 1.** Descriptive statistics and independent sample tests (according to age groups).

|  | Young<br>Mean (SD) | Old<br>Mean (SD) | t | p |
| --- | --- | --- | --- | --- |
| **Cognitive flexibility** |  |  |  |  |
| Rule Switching task | -7.17 (8.61) | -14.31 (9.22) | 3.31 | **.002** |
| Dogmatism Scale | 19.64 (4.28) | 20.01 (4.21) | -.38 | .71 |
| DOG | 40.21 (7.59) | 41.35 (8.91) | -.61 | .54 |
| Dependence on Routines | 19.02 (4.61) | 19.35 (5.15) | U=895<br>.51 | .61 |
| IRI-Perspective Taking | 26.31 (3.84) | 26.17 (3.72) | U=818<br>.51 | .84 |
| **Structural connectivity** |  |  |  |  |
| Global Efficiency | 5718.15 (257.92) | 5312.27 (563.99) | 3.85 | **<.001** |
| Local Efficiency | 582.53 (31.74) | 535.26 (64.94) | 3.85 | **<.001** |
| Clustering | 445.37 (26.93) | 412.08 (52.48) | 3.32 | **.002** |
| Strength | 126913.06 (4290.11) | 119511.42 (9767.81) | 4.08 | **<.001** |
| IFGpt clustering | 361.44 (40.69) | 311.83 (67.31) | 3.77 | **<.001** |
| IFGpt strength | 77364.7434 (12976.74) | 67869.83 (13542.47) | 3.06 | **.002** |
| IFGpt efficiency | 460.47 (53.23) | 397.59 (81.52) | 3.86 | **<.001** |
| IPL clustering | 638.48 (85.15) | 550.79 (128.89) | 3.39 | **.001** |
| IPL strength | 223743.08 (22208.59) | 203216.14 (33894.28) | 3.03 | **.004** |
| IPL efficiency | 822.06 (115.42) | 701.05 (165.27) | 3.65 | **<.001** |
| IIPL-lputamen | 5989.61 (2429.68) | 3518.41 (1916.76) | 4.84 | **<.001** |
| IIPL-rputamen | 350.17 (195.28) | 416.95 (296.41) | -1.14 | .26 |
| IIPL-lprecuneus | 3126.79 (2000.52) | 4041.27 (2441.21) | -1.75 | .083 |
| IIPL-rprecuneus | 2255.01 (1183.81) | 1204.51 (961.38) | 4.14 | **<.001** |

Ciphers in bold indicate values that survived multiple comparisons correction.

### 3.2 Relationship of cognitive flexibility and associated traits

The correlation matrices of cognitive flexibility with associated traits is displayed in Table 2. In young adults, a better performance in cognitive flexibility (Rule Switching task) was related to a lower dogmatism (DOG) and lower dependence on routines. In addition, a higher dogmatism (DOG) related to higher dependence on routines, and higher dogmatism (Dogmatism I) related to lower perspective taking. In older adults, a significant negative correlation was also observed between dogmatism (Dogmatism I) and perspective taking (i.e. higher levels of dogmatism were related to lower levels of perspective taking); no other significant correlation was observed in this group.

**Table 2.**
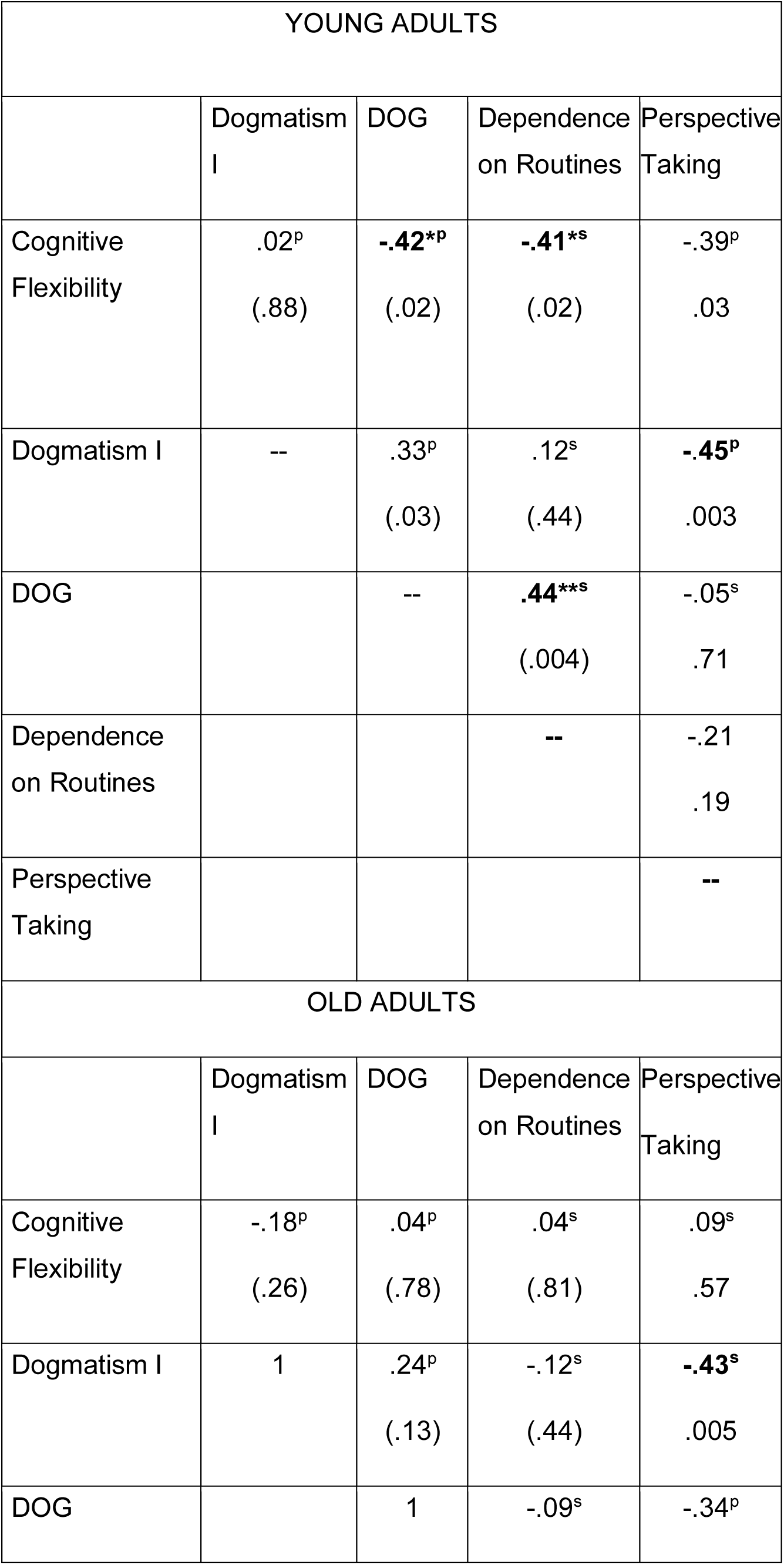
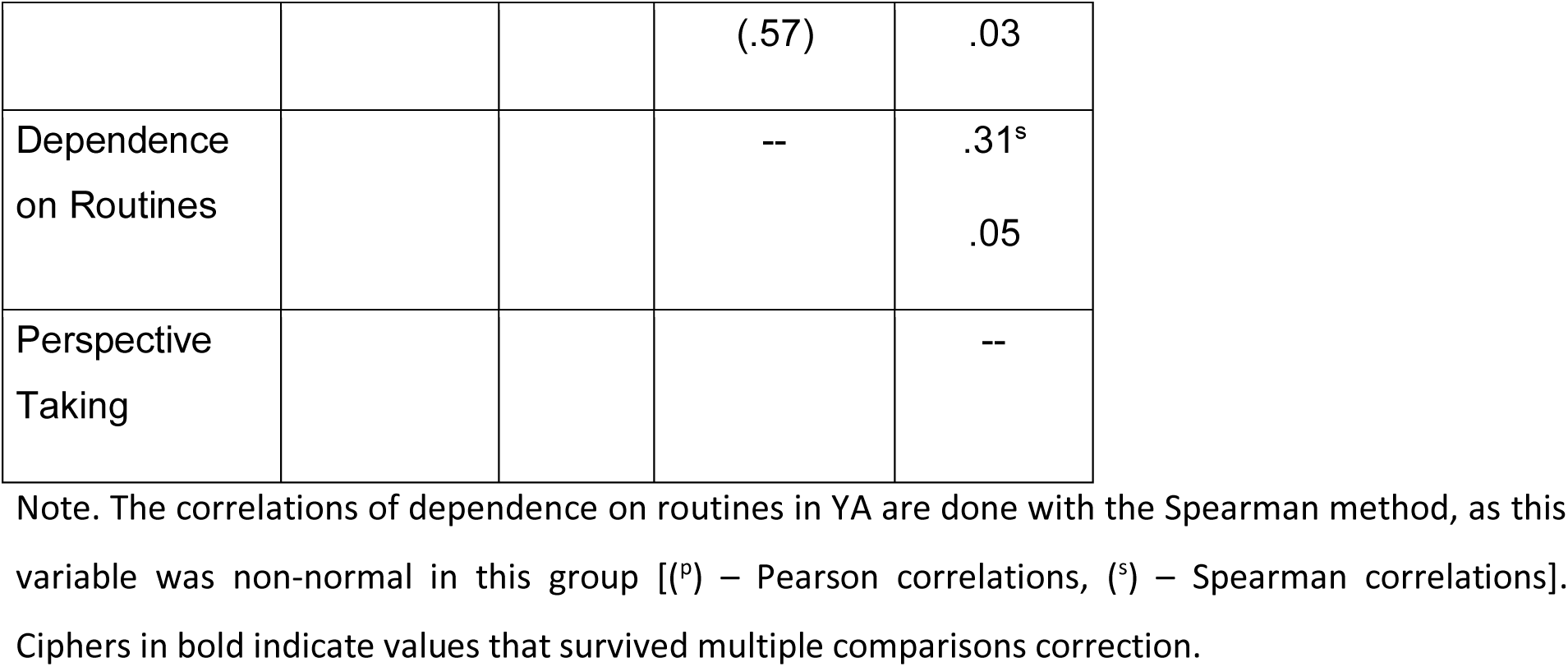
Correlations of Rule Switching with questionnaires.

### 3.3 Prediction of cognitive flexibility and associated traits from global structural connectivity properties

Regression (stepwise) models revealed that clustering negatively predicted dogmatism (Dogmatism I) in the full sample (i.e. a higher clustering was related to lower levels of dogmatism). In addition, strength negatively predicted dogmatism (DOG) in the full sample (Figure 3; Table 3): higher connection strength was related to lower dogmatism levels.

**Figure 3.**
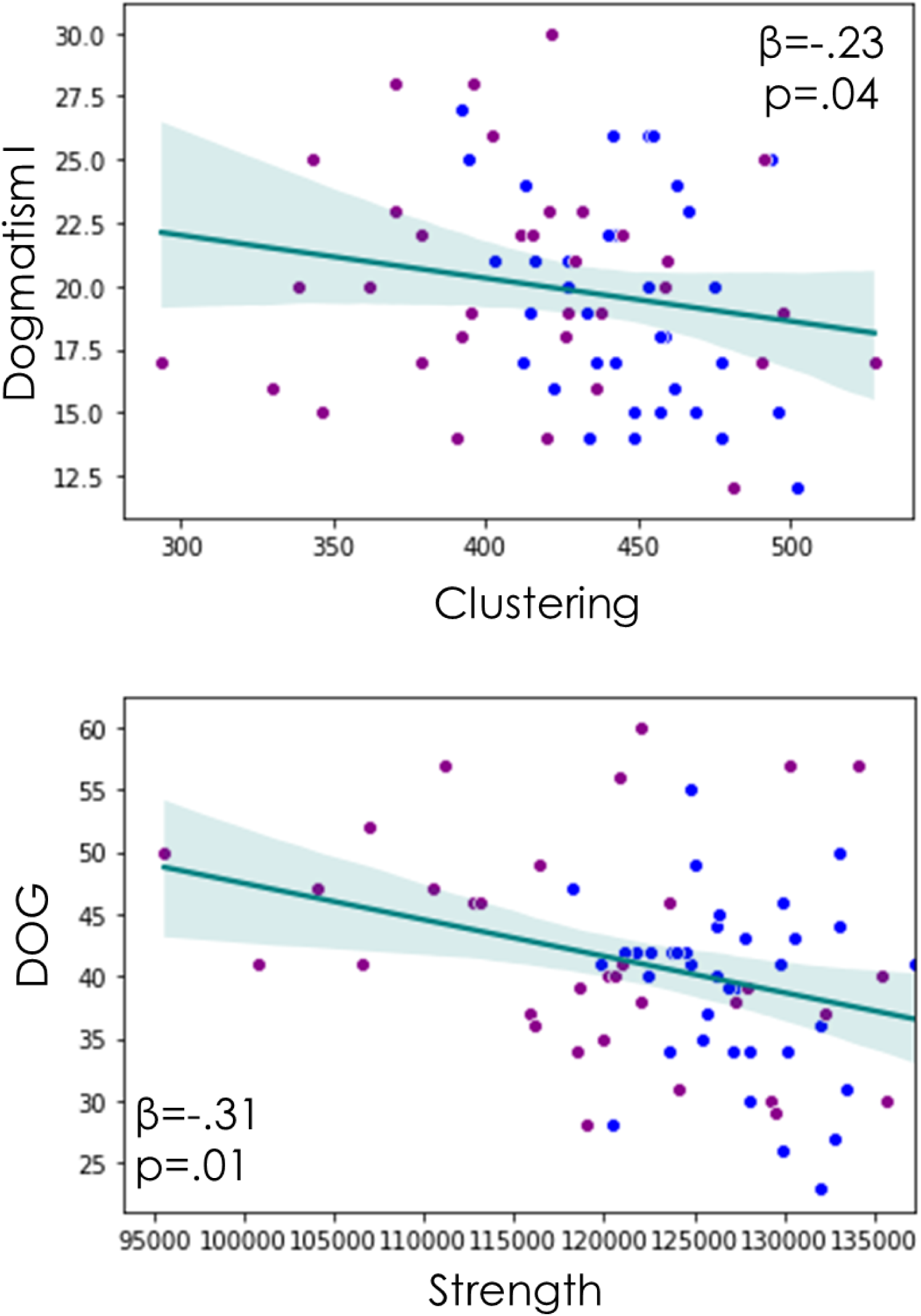
Prediction of cognitive flexibility associated traits from global connectivity properties. Top: Clustering negatively predicted dogmatism (Dogmatism I). Bottom: Connection Strength negatively predicted dogmatism (DOG). Blue dots – Young adults; Violet dots – Older adults.

**Table 3.** Regression models predicting cognitive flexibility and associated traits from global brain connectivity properties.

|  | Model |  |  | Predictors |  |  |  |
| --- | --- | --- | --- | --- | --- | --- | --- |
|  | R<sup>2</sup> | F (df<sub>reg</sub>, df<sub>tot</sub>) | p |  | β | t | p |
| *Rule Switching task* | .11 | 7.88 (1,63) | .007 | *Age* | -.33 | -2.81 | .007 |
| *Dogmatism I* | .06 | 4.28 (1,72) | .042 | *Clustering* | -.23 | -2.07 | .042 |
| *DOG scale* | .09 | 7.03 (1,70) | .01 | *Strength* | -.31 | -2.65 | .01 |
| *Dependence*<br>*on Routine* | -- | -- | -- | -- | -- | -- | -- |
| *Perspective*<br>*taking* | -- | -- | -- | -- | -- | -- | -- |

\*Age in years

### 3.4 Prediction of cognitive flexibility and associated traits from structural connectivity properties of the IFGpt and IPL

The connectivity strength in the full sample and the clustering from the same region in interaction with age significantly predicted the performance in the Rule Switching task. Posthoc analyses indicated that a higher clustering in IFGpt was related to a better cognitive flexibility performance in young adults but no significant relationship was observed in older adults (young: r=.52, p=.003; old: r=.36, p=.006; Figure 4; Table 4). The connectivity strength from IPL significantly and negatively predicted dogmatism (Dogmatism I; a higher connectivity strength was associated to lower dogmatism) in the full sample (Figure 5A, Table 4). Finally, the clustering of the IFGpt negatively predicted dogmatism (DOG; a higher clustering related to a lower dogmatism) in the full sample (Figure 5B; Table 4).

**Figure 4.**
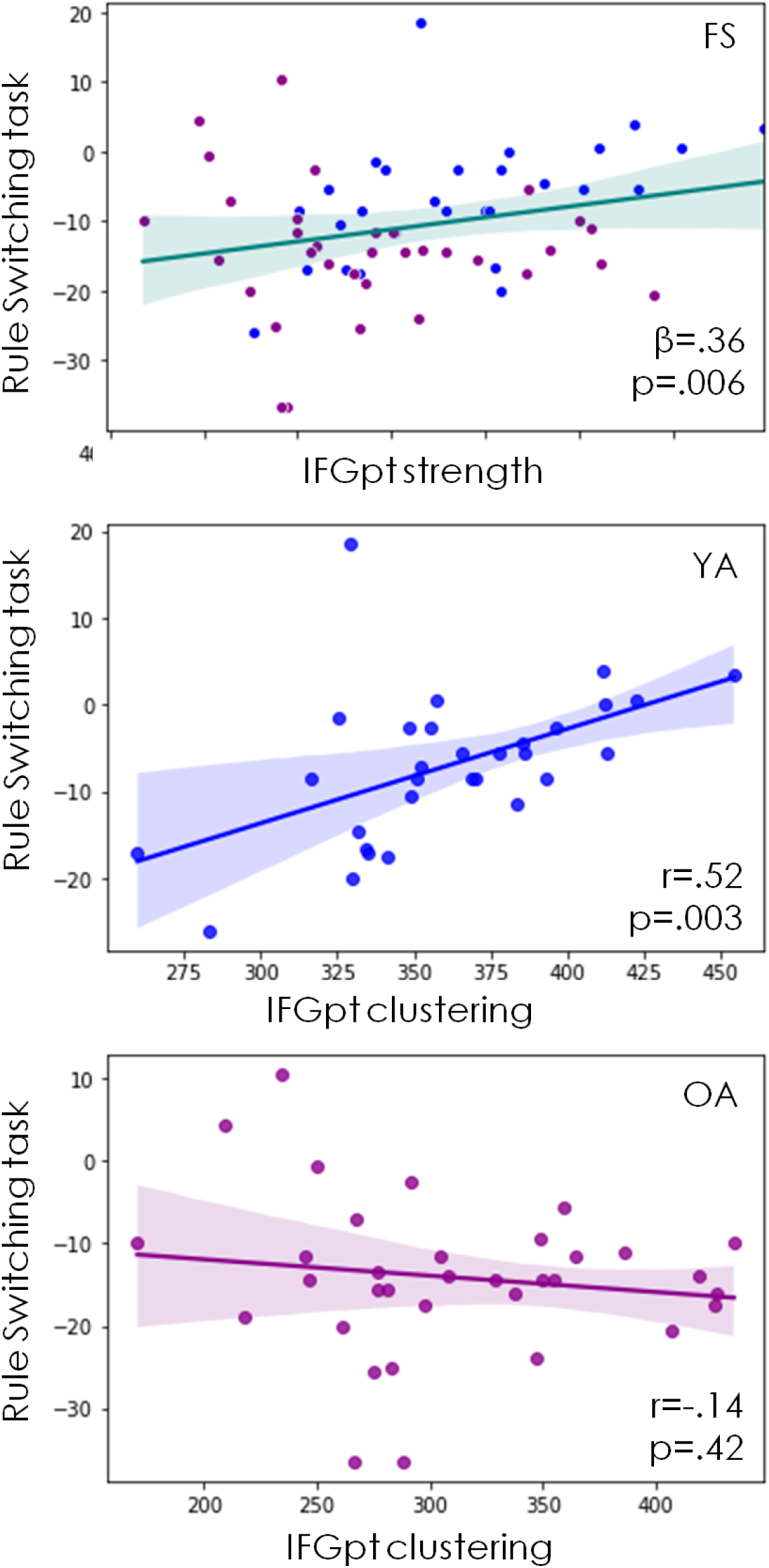
Prediction of cognitive flexibility from IFGpt connectivity properties. Top: The connectivity strength of the IFGpt predicted cognitive flexibility performance (Blue dots – Young adults; Violet dots – Older adults). Middle and Bottom: The clustering in the IFGpt predicted the cognitive flexibility performance in young adults (middle) but not in older adults (bottom). FS-full sample; YA-young adults; OA-older adults.

**Figure 5.**
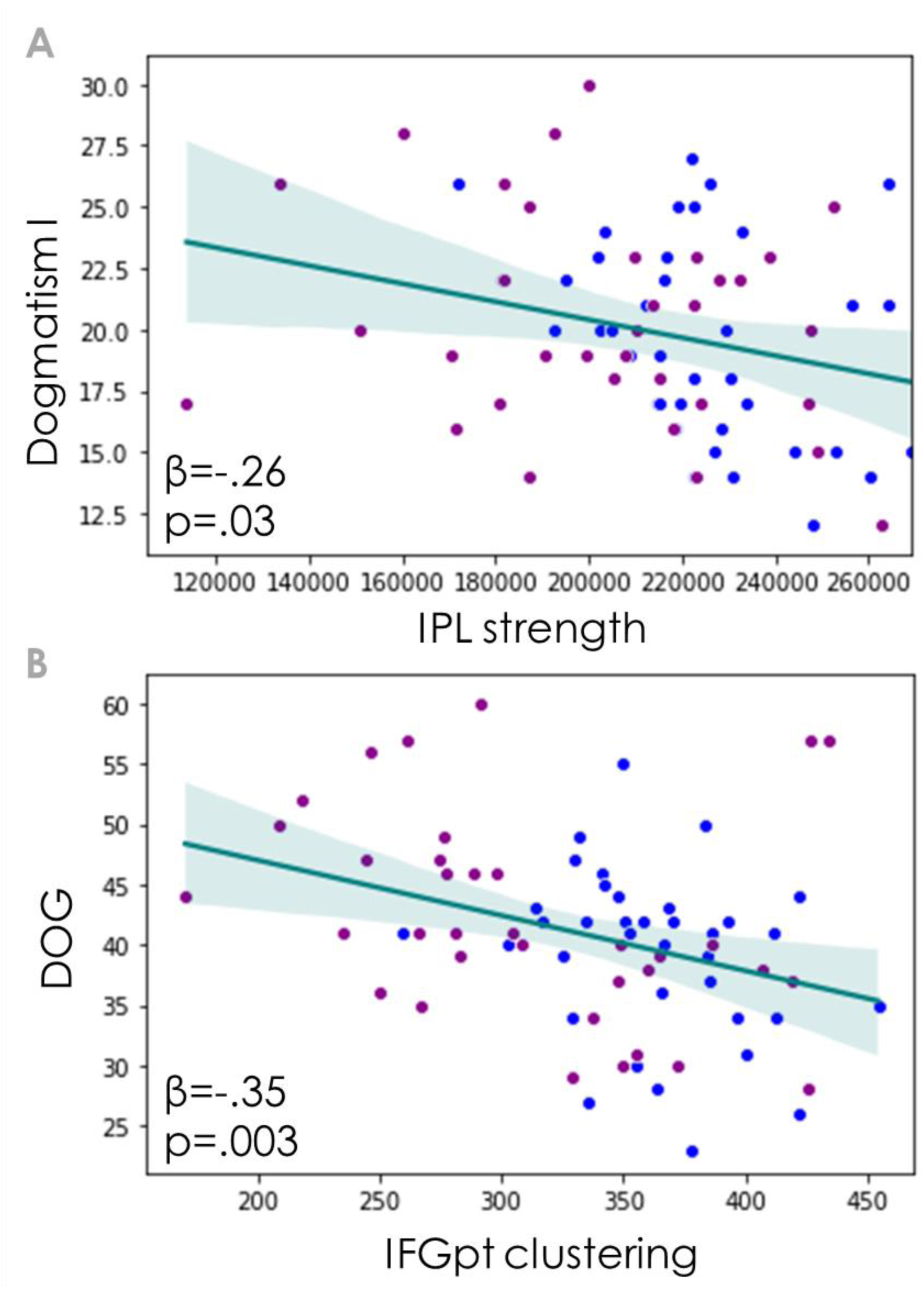
Prediction of cognitive flexibility associated traits from local connectivity properties. A) The connectivity strength in the IPL negatively predicted dogmatism (Dogmatism I) in the full sample. B) The clustering in the IFGpt negatively predicted dogmatism (DOG) in the full sample. Blue dots – Young adults; Violet dots – Older adults.

**Table 4.** Regression models predicting cognitive flexibility and associated traits from structural connectivity properties of the IFGpt and IPL.

|  | Model |  |  | Predictors |  |  |  |
| --- | --- | --- | --- | --- | --- | --- | --- |
|  | R<sup>2</sup> | F (df<sub>reg</sub>, df<sub>tot</sub>) | P |  | β | T | p |
| *Rule Switching task* | .14 | 5.07 (2,63) | .009 | *IFGpt strength* | .36 | 2.82 | .006 |
|  |  |  |  | *IFGpt clustering X Age* | -.29 | -2.36 | .021 |
| *Dogmatism Scale* | .07 | 5.17 (1,72) | .026 | *IPL strength* | -.26 | -2.27 | .026 |
| *DOG scale* | .17<br>.12 | 9.74 (1,70) | .001 | *IFGpt clustering* | -.35 | -3.12 | .003 |
| *Dependence on Routine* | -- | -- | -- | -- | -- | -- | -- |
| *Perspective taking* | -- | -- | -- | -- | -- | -- | -- |

### 3.5 Prediction of cognitive flexibility and associated traits from the IPL-putamen and IPL-precuneus connectivity strength

The connectivity strength between the left IPL and the left putamen in interaction with age negatively predicted dogmatism (DOG; Table 5). Posthoc analyses indicated that a stronger left IPL-left putamen connectivity was related to lower dogmatism (DOG) in older adults, but no relationship was observed in young adults (young: r=.18, p=.27 ; old: r=-.33, p = .05; Figure 6).

**Table 5.**
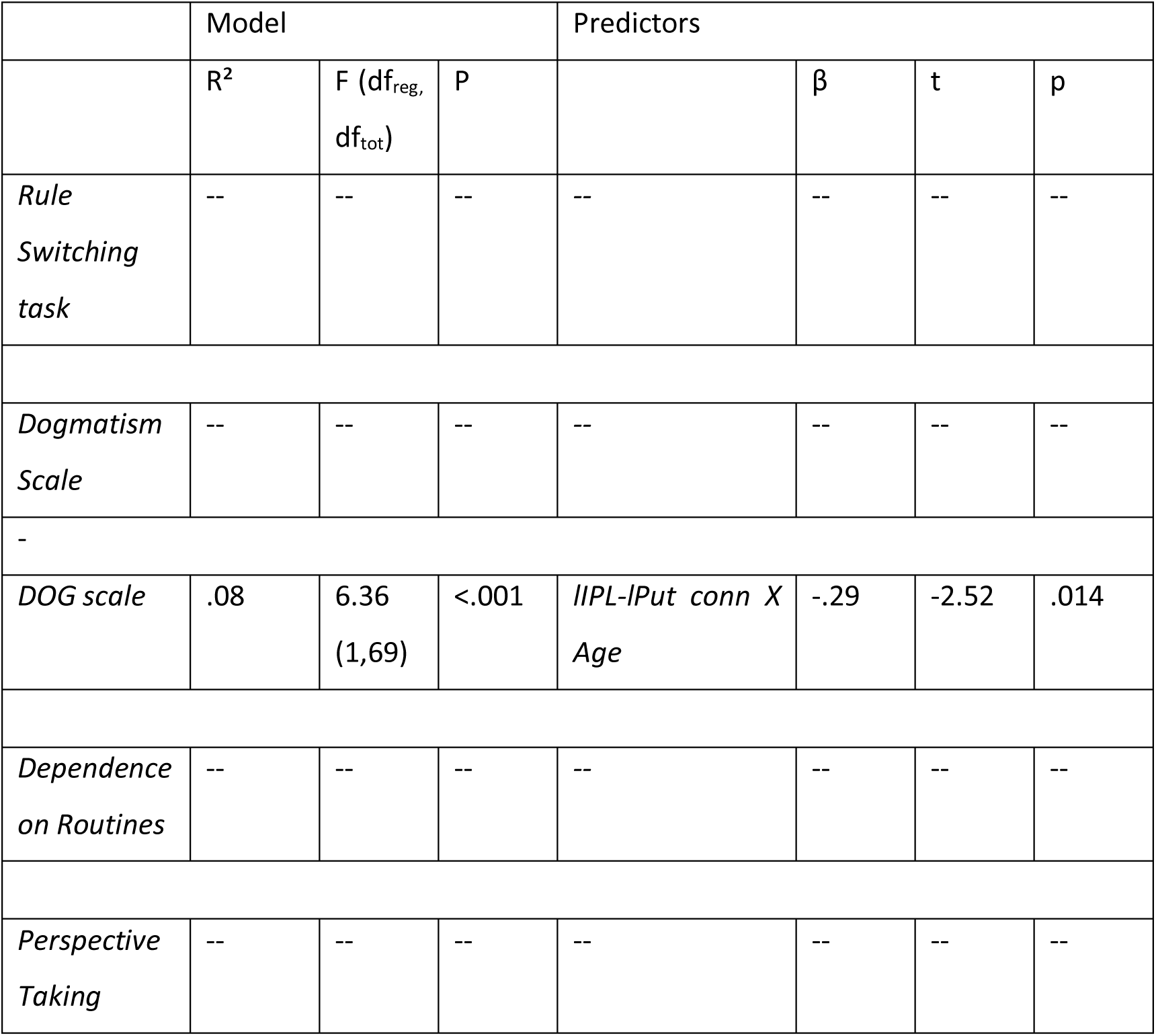
Regression models predicting cognitive flexibility and associated traits from structural connectivity of the left IPL with the bilateral putamen and the right precuneus.

**Figure 6.**
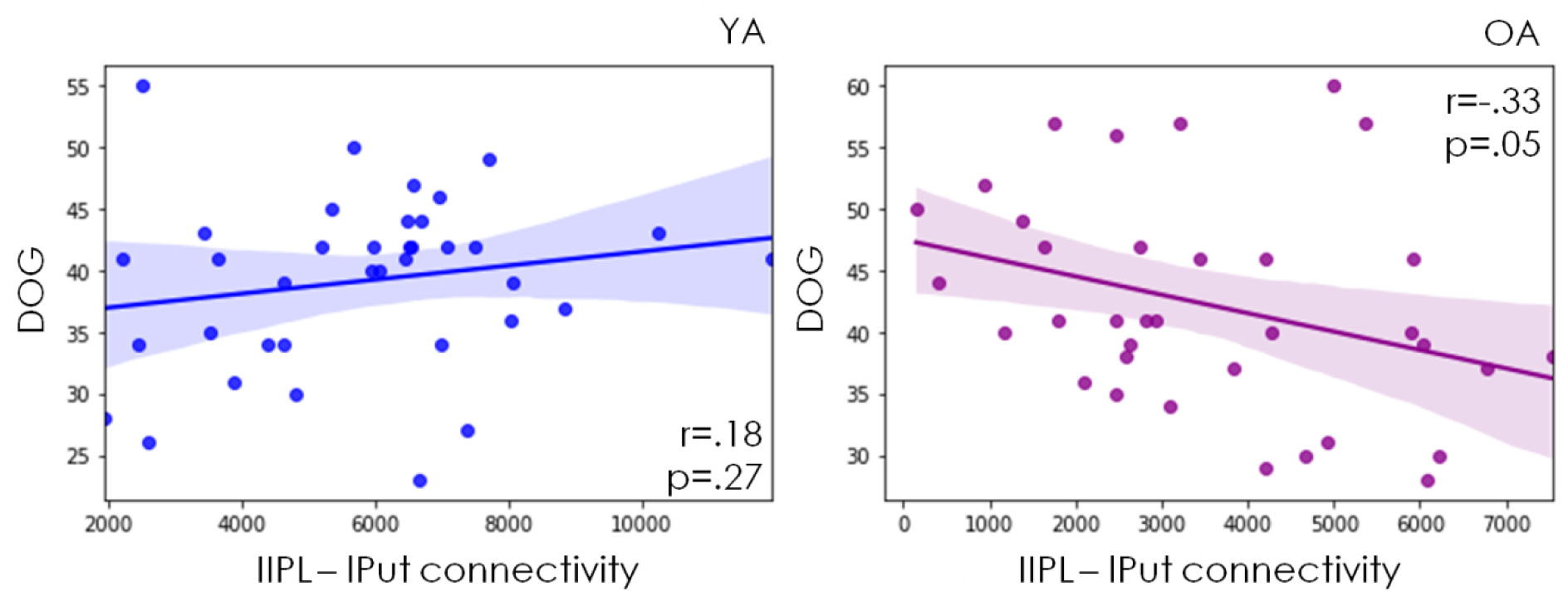
Prediction of cognitive flexibility associated traits from the connectivity of IPL. The left IPL – left Putamen connectivity strength was negatively associated with dogmatism (DOG) levels in older adults (right) but no relationship was observed in young adults (left).

## 4. DISCUSSION

We investigated the relationship between cognitive flexibility and associated traits (dogmatism, dependence on routines and perspective taking). We observed that a higher cognitive flexibility related to lower dogmatism and lower dependence on routines in young adults, but that relationship was absent in older adults. Importantly, there were no age differences in cognitive flexibility associated traits. Further, we investigated whether cognitive flexibility and associated traits could be predicted by different brain structural connectivity properties. We found that: a) global and local connectivity properties negatively predicted dogmatism levels in the full sample; b) local connectivity properties of the IFG positively predicted performance in cognitive flexibility performance in the full sample and in older adults, and c) the connectivity between lIPL and the left putamen negatively predicted dogmatism (DOG) in older adults. Next, we discuss these findings in detail.

### 4.1 Age differences in cognitive flexibility, associated traits and structural connectivity properties

In spite of older adults showing lower performance in the cognitive flexibility task, there were no age differences in either of the cognitive flexibility associated traits (dogmatism, dependence on routines and perspective taking). This finding contradicts the popular view that daily life cognitive flexibility associated traits are reduced with age. Although some bias may apply to this study, such as self-selection (possibly more active and open older adults are willing to engage in the study) and social desirability (answering in self-reports in a socially desirable manner), it can be argued that these biases may apply for the young individuals as well. It is also possible that neurodegenerative processes emerging with older age may increase the proneness to loose cognitive flexibility during daily tasks, but these neurodegenerative processes may not affect all older adults equally. Furthermore, it is important to mention that the cognitive flexibility associated traits may be related or considered to be part of stable personality traits (e.g. Openness to experience) which may be more resistant to age factors.

Regarding global connectivity properties, younger adults showed higher global and local efficiency, clustering and strength, replicating previous findings (Rodriguez-Nieto et al. 2025; Madden et al. 2020; Li et al. 2020). Likewise, younger adults showed higher indices of these properties in the IFG and in the IPL. These findings support a decreased structural connectivity in older adults that affects the efficiency of information transfer and a decreased segregation of information processing.

Interestingly, although younger adults showed a stronger connectivity between the left IPL and the left putamen, and between the lPL and right precuneus, there were no significant differences between young and older adults regarding the connectivity strength of the left IPL with the right putamen and the left precuneus. Whether these findings comprise compensatory mechanisms or are the result of a changes in cognitive processing as a result of alterations in life and cognitive style with aging remains to be elucidated.

### 4.2 Relationship of cognitive flexibility and associated traits

Whereas cognitive flexibility related to one dogmatism and the dependence on routines scales in young adults -revealing some degree of ecological validity of the Rule Switching task-this was not the case in older adults.

Although speculative, this observation could reflect that the Rule Switching task may not capture cognitive flexibility well in older people, perhaps as a result of altered working memory load.

### 4.3 Prediction of cognitive flexibility and associated traits from global structural connectivity properties

A higher degree of global clustering was related to lower levels of one scale of dogmatism (Shearman and Levine, 2006) in the full sample. Clustering refers to the identification of nodes that are densely connected to each other. In this sense, a higher clustering reveals a higher organizational complexity of information processing, which seems to relate to lower mental rigidity.

A higher degree of global connectivity strength was negatively related to the second dogmatism scale (Crowson et al. 2015) in the full sample. This is, a less densely connected global network related to a higher absolute certainty in one’s beliefs.

A richer and more complex global neural connectivity indicates a better communication among distinct and distant modules, which likely enables the capacity to connect different mental schemas, favouring a flexible way of thinking, a higher ability to take different perspectives to the same topic and to a more nuanced perspective into one’s beliefs which can benefit inter-personal understanding in daily life.

### 4.4 Prediction of cognitive flexibility and associated traits from structural connectivity properties of the IFGpt and IPL

The connectivity strength of the IFGpt was related to a better cognitive flexibility performance in the full sample, indicating that a stronger connectivity of this region predicts better skills in switching. The left IFGpt is a region that is engaged during the Rule Switching task paradigms (Rodriguez-Nieto et al., 2022). This finding shows that not only the local activation of this region but also its structural connectivity to other regions of the brain is relevant for successfully switching among mental schemes. In addition, the clustering of the IFGpt was related to better cognitive flexibility only in young adults, showing that, besides the connectivity strength, it is also the organizational complexity of that connectivity that appears to be relevant in young adults.

Interestingly, the connectivity properties of the two target hubs of the flexibility network (IFG and IPL) were distinctly related to specific forms of dogmatism. In particular, a higher clustering of the IFGpt was related to lower levels of Dogmatism (DOG; Crowson et al 2015) in the full sample, indicating that the connectivity complexity of this specific region is also relevant in evaluating the degree of absolute certainty of own’s beliefs. On the other hand, a higher connectivity strength of the IPL was related to lower dogmatism (as measured by the scale of Sherman and Levine, 2006), which indicates that a higher connectivity of this region relates to a higher openness to different approaches to the same topic or problem. To this regard, it is important to note that the IPL overlaps with the temporoparietal junction, a region that is crucial in social cognition and perspective taking (Igelström and Graziano, 2017).

### 4.5 Prediction of cognitive flexibility and associated traits from the IPL-putamen and IPL-precuneus connectivity

A higher connectivity strength of the left IPL with the left putamen predicted lower dogmatism (DOG; Crowson et al. 2015) in older individuals. This observation seems partially contradictory to our functional connectivity results, where we observed that a stronger lIPL-left putamen connectivity was related to poor cognitive flexibility performance only in older adults (Rodriguez Nieto et al. 2026). However, as noticed earlier, cognitive flexibility performance and dogmatism were not related in older individuals; thus functional and structural connectivity may just relate distinctively to different flexibility properties: stronger functional connectivity relating to lower cognitive flexibility performance (possibly reflecting a higher functional demand in individuals who struggle with the task) and stronger structural connectivity relating to lower dogmatism (reflecting the relevance of structural connectivity strength in a flexibility associated trait). Last, whereas previous literature has reported the role of fronto-striatal circuits in cognitive flexibility (Uddin 2021; Padron-Rivera 2022), the current evidence supports the view that parieto-striatal connectivity is also relevant for this executive functions.

### 4.6 Conclusion

This study showed that cognitive flexibility performance relates to individually associated traits in young adults but not in older adults. In addition, we showed that brain connectivity properties are associated to cognitive flexibility and different types of dogmatism. These results pave the way for new questions, such as whether it would be possible to facilitate (or reduce) the expression of certain traits through the training of a particular function and how plastic the structural networks are across the lifespan and their possible effect over the expression of particular traits.

Our observations are meaningful because they suggest that structural network properties (through their relation with cognitive flexibility and dogmatism) can substantially impact the adaptability to different environments, socio-emotional integration and social harmony. Importantly, we observed age-related differences in the relationship between brain connectivity features and cognitive flexibility and dogmatism. These results are relevant in view of the demographic evolution of society and the fast pace at which changes in the sociopolitical and environmental sphere occur.

## Acknowledgements

We deeply thank Aurélie Derom and Chrissy van de Voorde for their support in recruiting participants and during data collection. We also would like to extend our thankfulness to all participants for their cooperation.

